# Rats Exposed to a Low Resource Environment in Early Life Display Sex Differences in Blood Pressure, Autonomic Activity, and Brain and Kidney Pro-inflammatory Markers During Adulthood

**DOI:** 10.1101/2025.09.04.674369

**Authors:** Jonna Smith, Savanna Smith, Kylie Jones, Angie Castillo, Jessica L. Bolton, Ahfiya Howard, Luis Colon-Perez, Faith Femi-Ogunyemi, Allison Burkes, Mark Cunningham

## Abstract

**Introduction:** Poverty, a low resource state, is a common adverse childhood experience (ACE) and early life stress (ELS). People who experienced childhood poverty are at greater risk for developing hypertension during adulthood, with sex differences. To determine the possible mechanisms of these sex differences, we investigated the alterations in blood pressure (BP), autonomic activity, and inflammation in the brain and kidneys of rats exposed to an impoverished environment during the early postnatal period, by using the limited bedding and nesting (LBN) model. **Methods:** The LBN model mimics childhood poverty by creating a low resource environment on postnatal days 2-9. After weaning, offspring were separated by sex and LBN exposure and were evaluated at 16-18 weeks of age (Adulthood). **Results:** LBN males displayed an increase in BP compared to the control (CON), whereas LBN females showed no changes. Sympathetic nerve activity (SNA) was increased in LBN males and females compared to the CON, while only parasympathetic nerve activity (PNA) was increased in LBN vs. CON females. Pro-inflammatory cytokines, IL-17 and TNF-α, were decreased in the brains of LBN vs. CON males, with no alterations in females. **Conclusion:** Adult LBN males have elevated BP, due to increased SNA, while LBN females may be protected from increased BP due to a simultaneous increase in SNA and PNA. The reduction in IL-17 and TNF-α in LBN males may serve as a compensatory mechanism to lower BP. This study provides insights into sex differences in BP for adults who experienced childhood poverty.

## Introduction

Poverty impacts approximately 333 million children worldwide and increases the risk of developing hypertension and cerebrovascular diseases later in life^1^. Poverty is defined as resource deprivation that prevents individuals from maintaining a minimum standard of living^2^. Often, the definition of poverty is limited to its characterization of one’s economic status, such as a person with an income less than $30,000 a year in the United States (US)^3^. However, poverty is more than just a lack of income. Poverty restricts access to vital resources, such as food, water, shelter, transportation, education, jobs, safety, and healthcare^4^.

Poverty is a major public health concern that disproportionally affects children and is considered an adverse childhood experience (ACE). ACEs are traumatic events that occur before the age of 18 years, which can include neglect, abuse, household dysfunction, and importantly poverty^5–7^. According to the Centers for Disease Control (CDC), approximately 64% of the US population has experienced at least one type of ACE, while 17% reported experiencing four or more ACEs. ACEs can negatively impact a person’s mental and physical health throughout life^8^.

Epidemiological studies of ACEs and animal models of early life stress (ELS) report an increase in cardiovascular and cerebrovascular diseases during adulthood^9–13^. These studies not only show increased health risks during adulthood, but also sex differences in blood pressure (BP). Many studies show that males have a greater risk of developing hypertension, being diagnosed with hypertension at an earlier time point in adulthood, and display a larger increase in BP compared to females that have experienced an ELS^13–15^. However, these findings are controversial. In which some studies suggest no sex differences, while others show that post-menopausal women have equal or greater rates in hypertension compared to males who experienced an ACE^14^. The mechanisms that lead to these sex differences and increased risk of hypertension are unknown and the focus of this study.

Two mechanisms known to regulate BP and facilitate hypertension development are alterations in autonomic activity and inflammation within the brain and kidney^16–19^. Specifically, an increase in brain and kidney pro-inflammatory cytokines, such as interleukin-17 (IL-17) and TNF-alpha (TNF-α), are shown to increase BP in rodents with or without ELS, and with sex differences^18,20–22^. To study ELS in the context of a low resource environment, we utilized the limited bedding and nesting (LBN) rodent model to assess BP, autonomic activity, and alterations in inflammation within different regions of the brain and kidney.

The LBN model mimics childhood poverty by reducing bedding and nesting material during weaning^23,24^. This model creates a chronically stressed environment, characterized by unpredictable maternal care^25,26^. These fragmented behaviors observed in dams are also observed in people living in impoverished environments. In rodents, the radical changes in maternal care, such as alterations in duration and frequency of grooming, nest building, and rough handling, can mimic neglect and abuse experienced by children during poverty^24,26^. This ELS model has demonstrated that rodents exposed to LBN during the early postnatal period have alterations in brain structures (such as the hippocampus, amygdala, and prefrontal cortex), hypothalamic-pituitary-adrenal (HPA) axis, cognitive decline, and depression^26,27^. Despite these novel findings, no studies have investigated the sex differences and effect of LBN exposure on BP, autonomic activity, and inflammation in rodents during adulthood. These studies are necessary because ACEs are common in adults, and the physiological mechanisms connecting ACEs and increased risk of cardiovascular diseases and cerebrovascular dysfunction are unknown^13,28^.

The objective of this study is to characterize changes in BP, autonomic activity, and inflammation in the brain and kidney of male and female rats that experienced a low-resourced (LBN) environment during weaning. We hypothesize that LBN male rats will display elevated BP, changes in autonomic activity, and inflammation in the brain and kidney, whereas LBN females will display no changes in these measurements. The results from this study will provide insight into the mechanisms that link childhood poverty to the development of hypertension later in life, along with sex differences.

## Methods

### Experimental Procedure

Timed-pregnant Sprague-Dawley rats (ENVIGO; Indianapolis, IN) were received on gestational day (GD) 11-12 and were divided into two groups: Control (CON; n=6) and LBN (n=7) dams. The rats were housed in a 12-hr light/dark cycle with controlled temperature and humidity. Food and water were provided *ad libitum* throughout the entire experimental protocol. All animals and procedures used in this study were approved by the University of North Texas Health Fort Worth Institutional Animal Care and Use Committee (IACUC) and in accordance with the National Institute of Health (NIH) Guide for the Care and Use of Laboratory Animals.

At GD 20-22, both CON and LBN dams gave birth naturally. From postnatal day (PND) 1-21, the CON dams weaned their pups in a normal environment with the typical amounts of bedding and nesting material. From PND 2-9, LBN dams and pups were relocated to a LBN environment, with less bedding and nesting material, with 75-80% less bedding material to induce ELS^23,24,27^. The LBN environment included a slightly elevated mesh floor on the bottom of a clean cage to allow for droppings to fall through and to provide another layer of stress. During PND 2-6, behavior of both sets of dams were recorded, analyzed, and evaluated to calculate an entropy score, which is a measurement of fragmented behaviors for the dams^29,30^. The dams were monitored for frequency, total duration, and the mean duration of self-grooming, pup grooming, nursing, nest building, eating and drinking, and transportation of pups in the cage. All these measurements were observed to examine if there were any significant changes in behavior due to the LBN environment. The measurements were then used to generate an entropy score for our dams, using a computational algorithm^29,30^. The entropy score utilizes the conditional probability of each behavior, in sequence with other behavior types, to determine if the dams’ behavior was fragmented and unpredictable. Many other research groups have used the entropy score to validate the efficacy of the LBN model in generating ELS^23,24,29^. The wire mesh was removed, and normal quantities of bedding and nesting materials were provided to the LBN group on PND 10, which continued until PND 21. After the weaning period, all pups were separated by sex and treatment, then aged to 16-18 weeks of age, which is equivalent to young adulthood in humans^31^.

To measure the mean arterial pressure (MAP), heart rate (HR), sympathetic nerve activity (SNA), and parasympathetic nerve activity (PNA), carotid catheterization surgery was performed at 16-18 weeks of age. After the BP, HR, SNA, and PNA were recorded, we humanely euthanized the animals and collected blood and organs. The kidney and brain were snap-frozen and stored in a −80°C freezer. Later, the brains were sectioned into cerebrum, brainstem, and cerebellum, whereas the kidneys were separated into medulla and cortex for experimentation and analysis. The organ sections were then homogenized and centrifuged at 10,000G for 20 minutes to obtain supernatants for future use in colorimetric enzyme-linked immunosorbent assays (ELISAs), namely IL-17 and TNF-α. Whole blood was centrifuged at 2,000 revolutions per minute (RPM) for 10 minutes, to collect the plasma and measure corticosterone concentration.

### Carotid Catheterization Surgery

At 16-18 weeks of age, we performed carotid catheterization surgeries, as described previously by us and others^24,32,33^.

### BP, SNA, and PNA Recordings

The following day, systolic BP, diastolic BP, MAP, heart rate (HR), low frequency BP variability (LFBPV) for SNA, and high frequency HR variability (HFHRV) for PNA were recorded via a PowerLab 16/35 AD Instruments apparatus (ADInstruments, Australia) as performed and described previously^34^. To measure SNA, we examined the variability in MAP tracings at a low frequency output (0.2-0.8Hz) using power spectral analysis calculations^35^. To obtain PNA, we also used the power spectral analysis to analyze HR variability at the high frequency output (0.75-2.0Hz)^35^. The power spectral analysis values were calculated using the Fast Fourier Transform (FFT) algorithm (2,048 values, 50% overlapping segments)^35^.

### Inflammation Assays

To determine IL-17 and TNF-α concentrations in the brain and kidney, we performed the following ELISAs: IL-17 (DY8410) and TNF-α (DY510-05; R&D Systems, Minneapolis, MN) according to the manufacturer’s instructions and previous studies32. Before loading the samples into the 96-well plates, we diluted the kidney cortices to 1:10, while the kidney medullas were pipetted neat. The brain sections were also all pipetted without dilution. Both assays were read at 450nm using the BioTek Epoch 2 microplate reader (Agilent Technologies, Santa Clara, CA) alongside Generation 5 software (Santa Clara, CA).

### Corticosterone Assay

To evaluate stress, we measured plasma corticosterone concentrations using the Corticosterone ELISA kit (Cayman Chemical: Item No. 501320, Ann Arbor, MI)^24^. The assay was performed based on the manufacturer’s instructions with undiluted plasma.

### Statistics

To analyze dam behavior and entropy score, we utilized the Student’s t-test. Body weight, organ weight, BP, corticosterone, inflammation, and autonomic activity were analyzed by a two-way ANOVA with a Fisher’s least significant difference (LSD) post-hoc, using GraphPad Prism 10 (v. 10.0.3; Santa Clara, CA) and Excel (Microsoft). The data were reported as mean ± standard error of the mean (SEM), where statistical significance was defined as *P<0.05.

## Results

### LBN Induced Behavioral Changes and Increased Entropy Score in Dams

Dams exposed to the LBN model during PND 2-9 exhibited several behavioral modifications in frequency of events, total duration of events, and average time of each event during exposure to the LBN (**Table 1**). We observed increased frequency of licking and grooming of pups (10.2 ± 1.16 vs. 5.5 ± 1.07A.U.; P=0.03), self-grooming (7.74 ± 0.68 vs. 2.75 ± 0.64A.U.; P=0.0009), nest building (3.86 ± 0.84 vs. 1.9 ± 0.33A.U.; P=0.13), eating (4.23 ± 0.83 vs. 1.65 ±0.60A.U.; P=0.06), and off-nest (11.23 ± 1.14 vs. 5.5 ± 0.95 A.U.; P=0.008). Furthermore, the total duration of licking and grooming the pups (441.65 ± 52.94 vs. 283.80 ± 40.89s; P=0.07), self-grooming (258.83 ± 22.61 vs. 98.69 ± 30.34s; P=0.002), and eating (373.80 ± 80.49 vs. 137.73 ± 32.42s; P=0.06) also increased. However, there were no significant changes in total duration of nest building (144.11 ± 42.81 vs. 114.37 ± 29.20s; ns) and off-nest time (571.33 ± 150.59 vs. 405.15 ±178.83s; ns). There was a significant decrease in total duration of nursing time (1,803.63 ±150.85 vs. 2554.70 ± 166.52s; P=0.01) with LBN dams, but no change in nursing event frequency (9.89 ± 1.57 vs. 7.65 ± 0.82A.U.; ns). Overall, no significant changes in the average time of each behavior were observed during this period (**Table 1**). To validate the exposure of chronic stress on the dams and pups, an entropy score was calculated. The LBN dams showed increased entropy (1.16 ± 0.05 vs. 0.83 ± 0.02A.U.; P=0.001) compared to the CON. Other groups have also demonstrated a higher entropy score in LBN dams, reflecting a chronically stressed environment with fragmented maternal behaviors (**Table 1**)^23,24,27^.

**Table 1:**
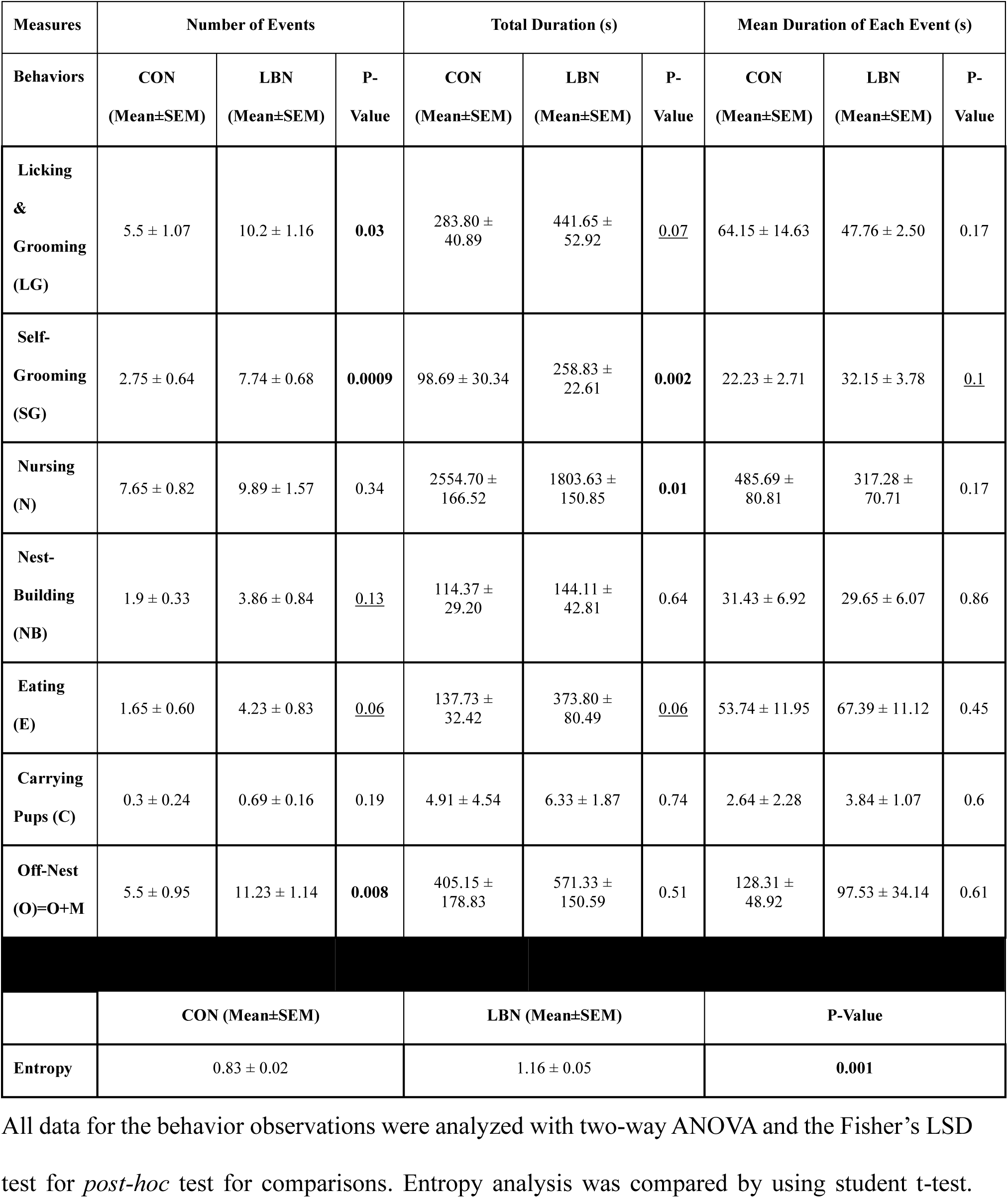
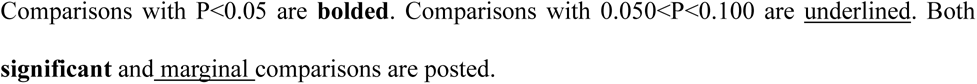
Behavioral Changes of Postpartum Dams (PND 2-6) Exposed to LBN.

| Measures | Number of Events |  |  | Total Duration (s) |  |  | Mean Duration of Each Event (s) |  |  |
| --- | --- | --- | --- | --- | --- | --- | --- | --- | --- |
| Behaviors | CON<br>(Mean±SEM) | LBN<br>(Mean±SEM) | P-Value | CON<br>(Mean±SEM) | LBN<br>(Mean±SEM) | P-Value | CON<br>(Mean±SEM) | LBN<br>(Mean±SEM) | P-Value |
| Licking<br>&<br>Grooming<br>(LG) | 5.5 ± 1.07 | 10.2 ± 1.16 | **0.03** | 283.80 ±<br>40.89 | 441.65 ±<br>52.92 | <u>0.07</u> | 64.15 ± 14.63 | 47.76 ± 2.50 | 0.17 |
| Self-Grooming<br>(SG) | 2.75 ± 0.64 | 7.74 ± 0.68 | **0.0009** | 98.69 ± 30.34 | 258.83 ±<br>22.61 | **0.002** | 22.23 ± 2.71 | 32.15 ± 3.78 | <u>0.1</u> |
| Nursing<br>(N) | 7.65 ± 0.82 | 9.89 ± 1.57 | 0.34 | 2554.70 ±<br>166.52 | 1803.63 ±<br>150.85 | **0.01** | 485.69 ±<br>80.81 | 317.28 ±<br>70.71 | 0.17 |
| Nest-Building<br>(NB) | 1.9 ± 0.33 | 3.86 ± 0.84 | <u>0.13</u> | 114.37 ±<br>29.20 | 144.11 ±<br>42.81 | 0.64 | 31.43 ± 6.92 | 29.65 ± 6.07 | 0.86 |
| Eating<br>(E) | 1.65 ± 0.60 | 4.23 ± 0.83 | <u>0.06</u> | 137.73 ±<br>32.42 | 373.80 ±<br>80.49 | <u>0.06</u> | 53.74 ± 11.95 | 67.39 ± 11.12 | 0.45 |
| Carrying<br>Pups (C) | 0.3 ± 0.24 | 0.69 ± 0.16 | 0.19 | 4.91 ± 4.54 | 6.33 ± 1.87 | 0.74 | 2.64 ± 2.28 | 3.84 ± 1.07 | 0.6 |
| Off-Nest<br>(O)=O+M | 5.5 ± 0.95 | 11.23 ± 1.14 | **0.008** | 405.15 ±<br>178.83 | 571.33 ±<br>150.59 | 0.51 | 128.31 ±<br>48.92 | 97.53 ± 34.14 | 0.61 |
|  | CON (Mean±SEM) |  |  | LBN (Mean±SEM) |  |  | P-Value |  |  |
| Entropy | 0.83 ± 0.02 |  |  | 1.16 ± 0.05 |  |  | **0.001** |  |  |

All data for the behavior observations were analyzed with two-way ANOVA and the Fisher's LSD test for *post-hoc* test for comparisons. Entropy analysis was compared by using student t-test.
Comparisons with $P < 0.05$ are **bolded**. Comparisons with $0.050 < P < 0.100$ are underlined. Both **significant** and marginal comparisons are posted.

### Corticosterone, Body Weight, and Organ Weight Analysis

Corticosterone, as a marker of stress, was measured in the plasma of adult offspring that experienced the LBN model (**Table 2**). At 16-18 weeks of age, there were no treatment effects or interaction effect for corticosterone in LBN exposed adults. However, there was a sex effect, showing that male rats had higher corticosterone concentrations compared to female rats (P=0.0034). Body weight, brain weight, total kidney weight, and heart weight of the of adult offspring showed no differences between treatment in male or female LBN vs. CON rats (**Table 2**). However, there was a sex effect, with males having a larger body weight and smaller brain-to-body weight ratio than females in both LBN and CON rats (**Table 2**).

**Table 2:** Measurement of Corticosterone, Body Weight, and Organ Weights.

|  | CON M | LBN M | CON F | LBN F | Treatment Effect (P-Value) | Sex Effect (P-Value) | Interaction (P-Value) |
| --- | --- | --- | --- | --- | --- | --- | --- |
| | (AVG ± SEM) | (AVG ± SEM) | (AVG ± SEM) | (AVG ± SEM) | | | |
| **Corticosterone (pg/mL)** | 4.28 ± 0.04 | 4.27 ± 0.11 | 4.07 ± 0.04 | 3.91 ± 0.04 | 0.3044 | **0.0034** | 0.366 |
| **Body Weight (g)** | 446.13 ± 13.28 | 444.43 ± 12.01 | 256.00 ± 6.71 | 259.25 ± 5.92 | >0.9999 | **<0.0001** | 0.7369 |
| **Brain Weight (g/kg of BW)** | 4.45 ± 0.12 | 4.44 ± 0.07 | 7.18 ± 0.17 | 6.90 ± 0.18 | 0.2933 | **<0.0001** | 0.3902 |
| **Total Kidney Weight (g/kg of BW)** | 5.64 ± 0.09 | 5.73 ± 0.14 | 5.81 ± 0.14 | 5.66 ± 0.19 | 0.8785 | 0.7696 | 0.3981 |
| **Heart Weight (g/kg of BW)** | 3.00 ± 0.09 | 3.21 ± 0.13 | 3.24 ± 0.19 | 3.25 ± 0.23 | 0.6408 | 0.2100 | 0.6785 |

All measurements at 16-18 weeks (adulthood) were analyzed with two-way ANOVA and the Fisher's LSD test for *post-hoc* test for comparisons. Comparisons with $P < 0.05$ are **bolded**.

### LBN & Sex: Effects on BP, HR, and Autonomic Activity

Systolic BP, diastolic BP, MAP, HR, and autonomic activity (SNA and PNA) were measured in adult CON and LBN rats (**Figures 1A-D and 2A-B)**. Both systolic (**Figure 1A**) and diastolic BP (**Figure 1B**) did not show an overall LBN exposure or sex effect. However, sex and LBN status did show a tendency for an interaction (Systolic BP: P=0.10; Diastolic BP: P=0.07) effect (**Figures 1A-B**). When checking for multiple comparisons, LBN males had elevated systolic (141 ± 6 vs. 122 ± 7mmHg, P=0.06) (**Figure 1A**) and diastolic (132 ± 5 vs. 114 ± 5mmHg, P=0.03) (**Figure 1B**) BP compared to CON males, whereas females no differences. Moreover, MAP showed both a treatment and interaction effect, in which LBN exposure had an effect in males but no effect in females (**Figure 1C**). In fact, LBN males exhibited a 16% (∼23mmHg) increase in MAP (139 ± 3 vs. 117± 5mmHg; P=0.0001) compared to the CON males (**Figure 1C**). There were no changes in MAP between LBN and CON females. There were no differences in HR among the groups (**Figure 1D**).

**Figure 1.**
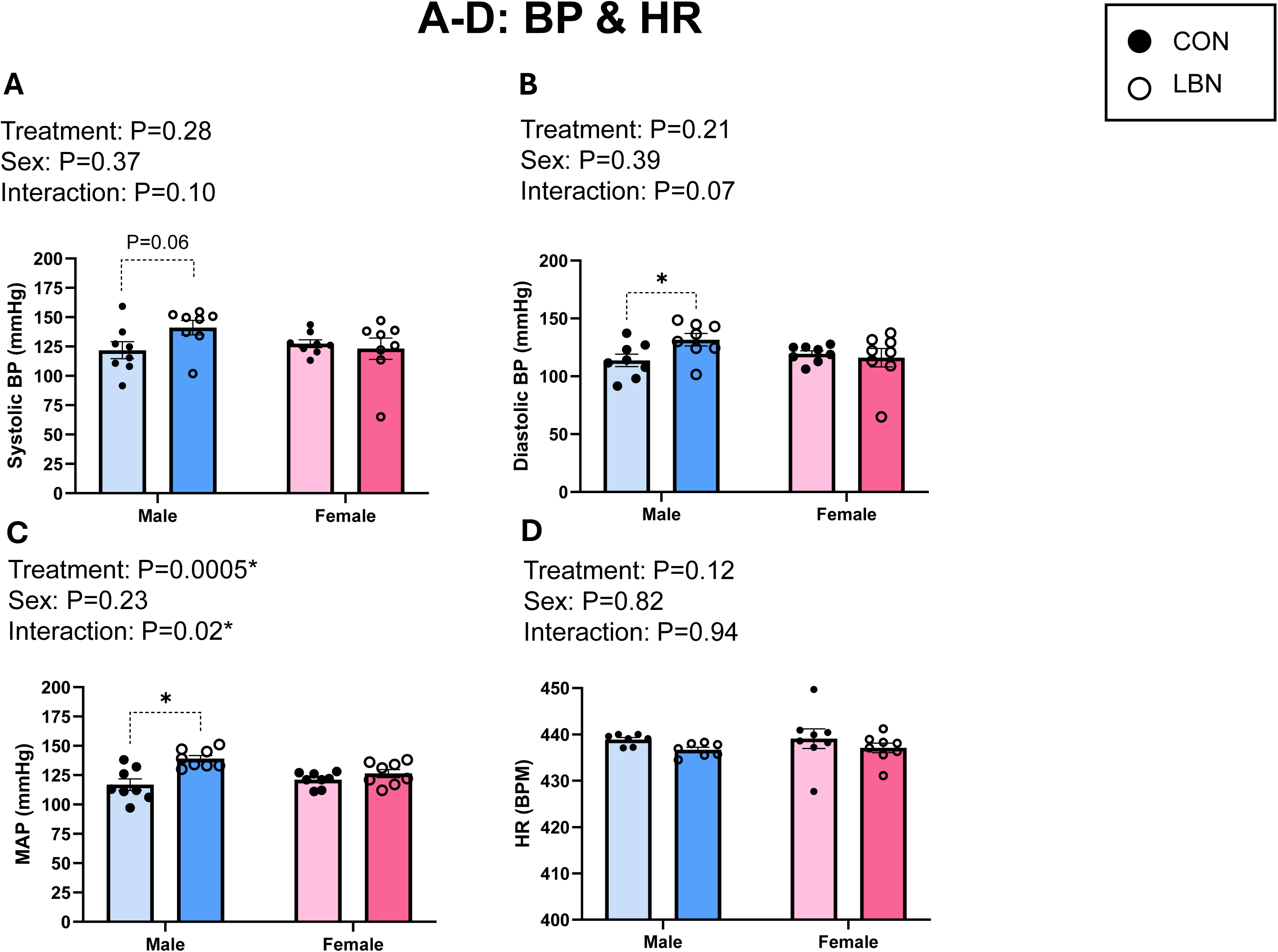
**A**) Systolic BP, **B)** diastolic BP, **C)** MAP, and **D)** HR were measured in LBN males (*n*=7-8), CON males (*n*=7-8), LBN females (*n*=8), and CON females (*n*=8). All bar graphs display individual data points with each group’s mean ± SEM, with significance determined at *P*<0.05. Groups were statistically analyzed with two-way ANOVA and the Fisher’s LSD test for *post-hoc* analysis. Individual data points are represented by closed circles for CON offspring and open circles for LBN offspring. Male bar graphs are blue and female bar graphs are pink.

To determine autonomic activity, we measured LFBPV (SNA) and HFHRV (PNA). There was a significant increase in SNA with LBN exposure in both males and females (P=0.0077) (**Figure 2A**). LBN males had a 6-fold increase in SNA (0.218 ± 0.07 vs. 0.04 ± 0.02mmHg^2^, P= 0.06) and LBN females displayed a 23-fold increase in SNA (0.209 ± 0.10 vs. 0.009 ± 0.01mmHg^2^, P=0.05) compared to their respective CON groups (**Figure 2A**). Conversely, PNA was only increased in LBN females vs. CON females (10,199.71 ± 2,047.22 vs. 5,004.83 ± 892.29ms^2^; P=0.01) (**Figure 2B**). No changes were observed in PNA between LBN males and CON males (ns).

**Figure 2.**
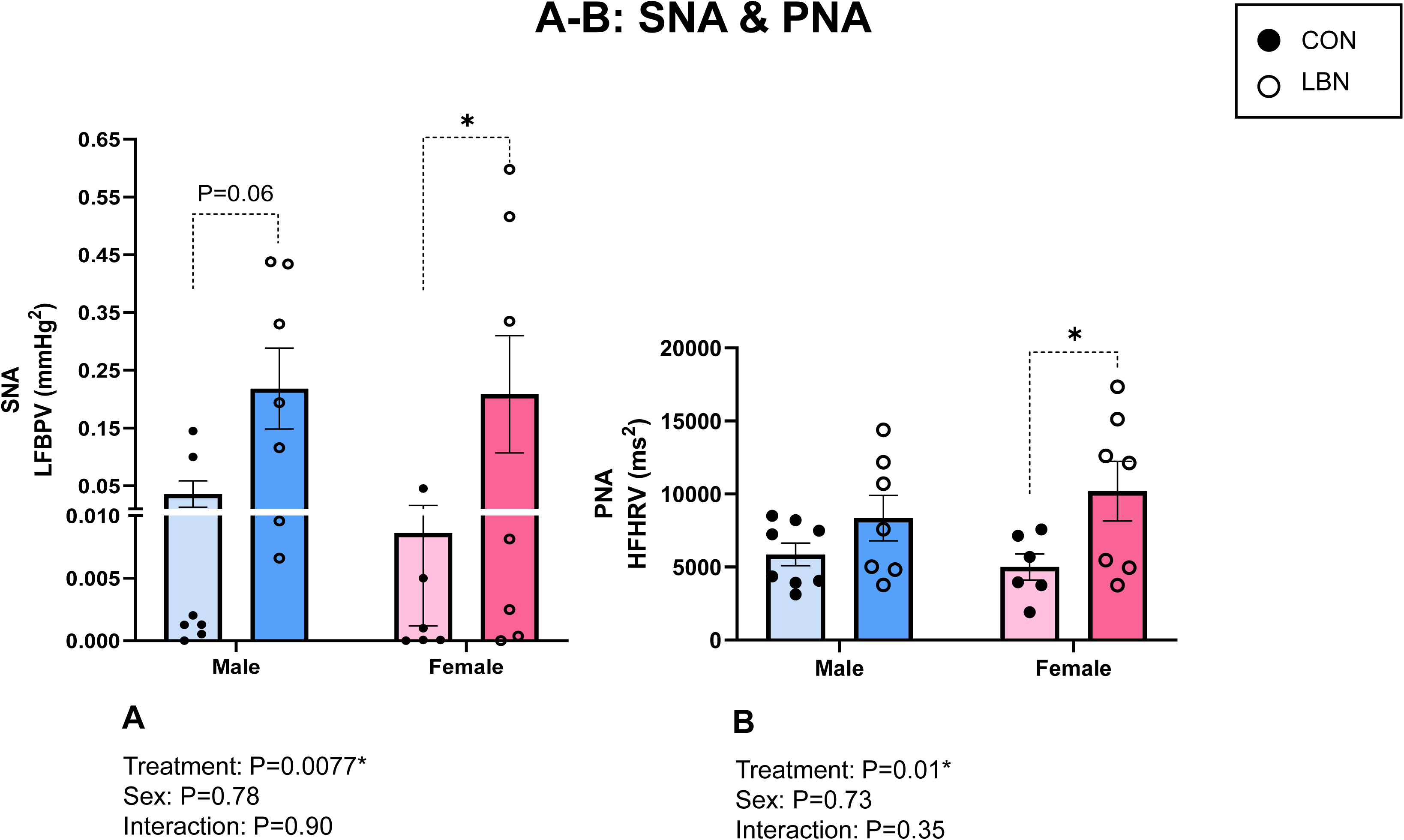
**A)** Low Frequency BP Variability (LFBPV; SNA) and **B)** High Frequency Heart Rate Variability (HFHRV; PNA) were measured in LBN males (n=7), CON males (n=7-8), LBN females (n=7), and CON females (n=6). All bar graphs display individual data points with each group’s mean ± SEM, with significance determined at *P*<0.05. Groups were compared with two-way ANOVA and the Fisher’s LSD test for *post-hoc* analysis. Individual data points are represented by closed circles for CON offspring and open circles for LBN offspring. Male bar graphs are blue and female bar graphs are pink.

### LBN & Sex: Effects on Brain Inflammation

IL-17 and TNF-α were examined in different segments of the brain: cerebrum, brainstem, and cerebellum outlined in **Figures 3A-F**. There was a trend in the interaction effect (sex X LBN), observed in the cerebrum (P=0.08) and brainstem (P=0.06) for IL-17. There were no significant changes in cerebral IL-17 in females (**Figure 3A**). Although in **Figures 3A-B**, LBN males exhibited a decrease in IL-17 concentrations in the cerebrum (1.93 ± 0.13 vs. 2.96 ± 0.34mg/mL/mg of Protein, P=0.01) and brainstem (2.04 ± 0.35 vs. 4.43 ± 4.43 ± 1.23mg/mL/mg of Protein; P=0.02). There was no difference in cerebellar IL-17 between LBN vs. CON rats within each sex, but there was an overall sex difference (P=0.02) with females having less IL-17 compared to males (**Figure 3C**).

**Figure 3.**
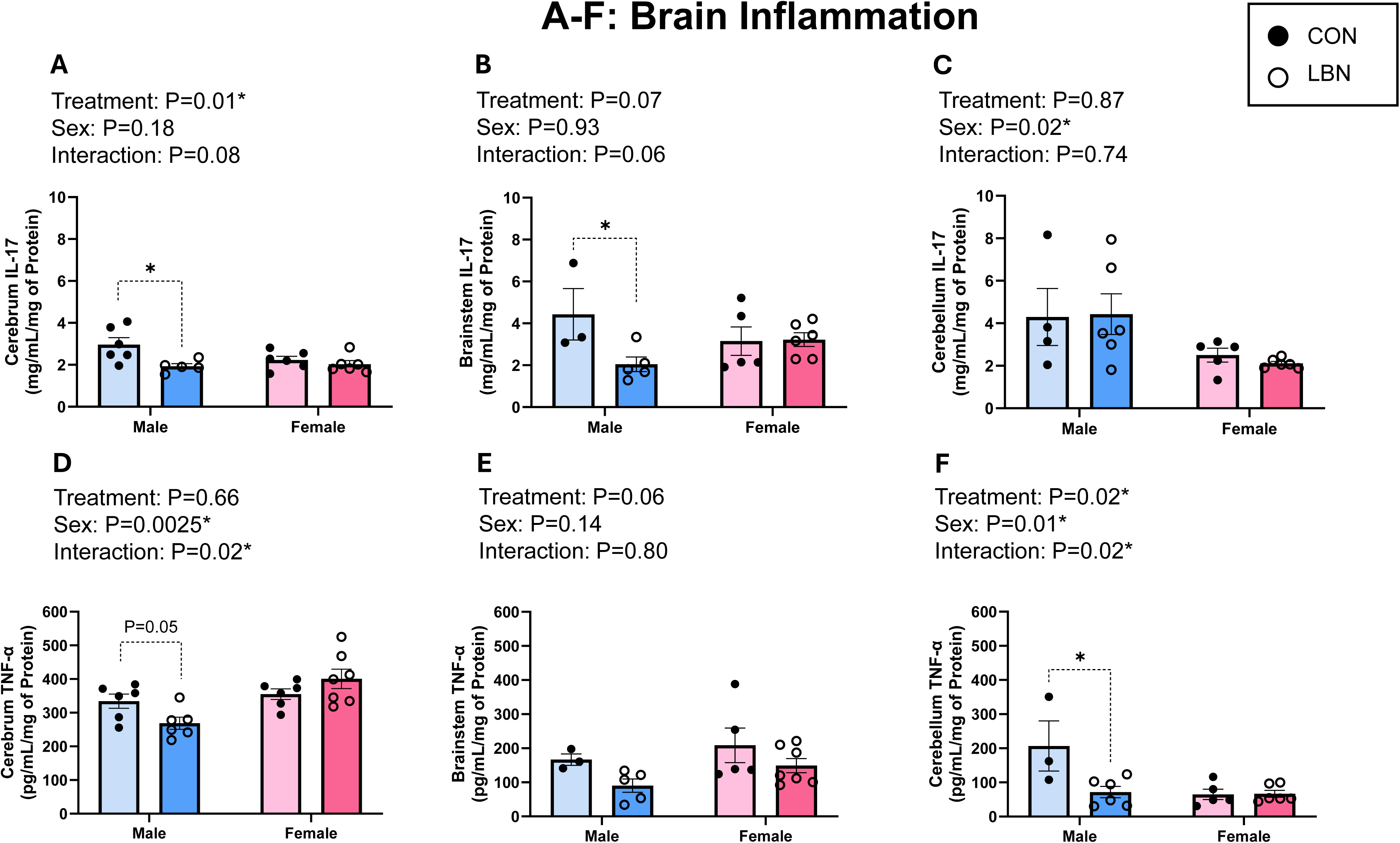
**A)** Cerebrum, **B)** brainstem, and **C)** cerebellum IL-17 concentrations were measured in LBN males (n=5-6), CON males (n=3-6), LBN females (n=5-6), and CON females (n=6). **D)** Cerebrum, **E)** brainstem, and **F)** cerebellum TNF-α concentrations were measured in LBN males (n=5-6), CON males (n=3-6), LBN females (n=6-7), and CON females (n=5-6). All bar graphs display individual data points with each group’s mean ± SEM, with significance determined at *P*<0.05. Groups were compared with two-way ANOVA and the Fisher’s LSD test for *post-hoc* analysis. Individual data points are represented by closed circles for CON offspring and open circles for LBN offspring. Male bar graphs are blue and female bar graphs are pink.

Cerebral TNF-α concentration was decreased for LBN vs. CON males (268.81 ± 17.99 vs. 334.28 ± 20.91pg/mL/mg of Protein; P=0.05), with no differences in LBN vs. CON females (**Figure 3D**). No differences were observed with brainstem TNF-α (**Figure 3E**). However, cerebellum TNF-α showed a 4-fold decrease between LBN vs. CON males (71.90 ± 16.43 vs. 206.79 ± 73.65pg/mL/mg of Protein; P=0.003), with no changes presented in females (**Figure 3F**).

### LBN & Sex: Effects on Kidney Inflammation

Concentrations of IL-17 and TNF-α were also investigated within the kidney cortex (**Figures 4A & C)** and medulla (**Figures 4B & D)**. IL-17 within the kidney cortex revealed no effects with LBN exposure and/or sex (**Figure 4A**). However, IL-17 in the kidney medulla tended to decrease in LBN vs. CON males (4.55 ± 0.33 vs. 5.83 ± 0.64mg/mL/mg of Protein; P=0.08) (**Figure 4B**). No differences exist between sex and LBN exposure vs. CON females for kidney medulla IL-17.

**Figure 4.**
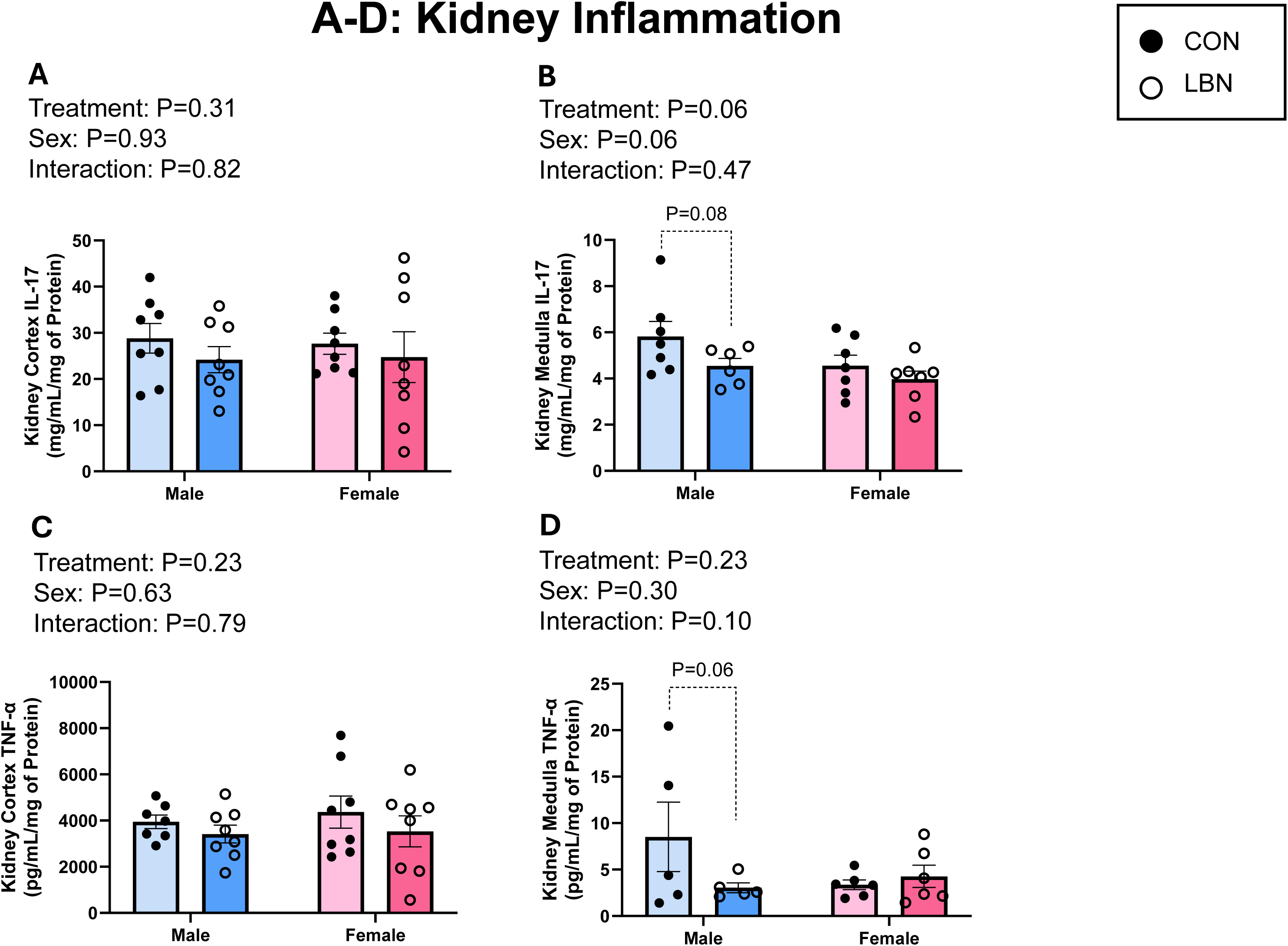
**A)** Kidney cortex and **B)** kidney medulla IL-17 concentrations were measured in LBN males (n=6-8), CON males (n=7-8), LBN females (n=7-8), and CON females (n=7-8). **C)** Kidney cortex and **D)** medulla TNF-α concentrations were also measured in LBN males (n=5-8), CON males (n=5-7), LBN females (n=6-8), and CON females (n=6-8). All bar graphs display individual data points with each group’s mean ± SEM, with significance determined at *P*<0.05. Groups were compared with two-way ANOVA and the Fisher’s LSD test for *post-hoc* analysis. Individual data points are represented by closed circles for CON offspring and open circles for LBN offspring. Male bar graphs are blue and female bar graphs are pink.

In the kidney cortex, there was no differences in LBN exposure and/or sex between groups for TNF-α (**Figure 4C**). Although, kidney medulla TNF-α tended to decrease in LBN vs. CON males (3.04 ± 0.53 vs. 8.52 ± 3.74pg/mL/mg of Protein; P=0.06) (**Figure 4D**).

## Discussion

LBN exposure induces sex differences in BP, autonomic activity, and inflammation in the brain and kidney. Specifically, LBN males exhibit an increase in BP, increased SNA, and decreased pro-inflammatory markers in the cerebrum, brainstem, and cerebellum. LBN females depicted no differences in BP, an increase in both SNA and PNA, and no alterations in brain or kidney inflammation. Results from our study suggests that the elevation in BP for LBN males may be due to an increase in SNA. However, LBN females may be protected against BP elevation due to a simultaneous increase in PNA, despite increased SNA. The sex differences in inflammation may be a compensatory response to BP changes, in which LBN males show a reduction in inflammation to mitigate brain damage from an increase in BP. Meanwhile, LBN females do not show changes in inflammation, since BP is not different between groups.

Children exposed to ACEs, including poverty, have an increased risk of poor health outcomes such as a heightened risk of hypertension, inflammation, and stroke as adults^13,36^. In addition, several clinical studies show that ACEs are linked to sex differences in the presentation of hypertension^7,37–40^. For example, in Huang *et al.’s* study, they found that persons experiencing ACEs increased the risk of hypertension later in life, indirectly, through cardiometabolic dysregulations (such as hyperlipidemia and hyperglycemia), systemic inflammation, and obesity^37^. In a similar study, conducted in the United Kingdom, Deschenes *et al.* found that British civil service employees, who reported ACEs had a higher risk for developing heart coronary disease, partially mediated through cardiometabolic dysregulation and hypertension^39^. Additionally, Su *et al.’s* 2015 longitudinal study, there was a strong, positive relationship between the number of ACES and BP, in which participants who were exposed to more ACEs exhibited a greater increase in BP during early adulthood compared to those who did not experience any ACEs^7^.

The association between elevated BP and ACEs is not only observed in human studies, but also in animal models of ELS, such as the maternal separation and/or early weaning models, which are animals models with shortened weaning periods^41–43^. Franco *et al.’s* 2013 study revealed that 25-week-old adult Winstar male rats exposed to early weaning were hypertensive with extensive oxidative stress^41^. In another study, Reho and Fisher 2015 investigated changes in vascular function and BP in a maternal separation rodent model. In these studies, they found no changes in BP, but an increase in arterial contractility at PND 21. However, at PND 35, the relationship was reversed, in which BP was increased while arterial contractility was unchanged^42^. Furthermore, in a study performed by Genest *et al.* 2004, showed that neonatal maternal separation led to a stark 20% increase in MAP in males, but no changes in females at 8-10 weeks of age^43^. Note that these results are similar to our findings, in which LBN males, at 16-18 weeks of age, had significantly increased MAP, while females remained unchanged. Together, these data show that ELS, whether it be via maternal separation or our model of LBN, is linked to hypertension.

The LBN ELS model is a resource deprivation model that mimics chronic stress experienced by the dams and pups during weaning. Typically, the LBN model is used to evaluate outcomes of depression and cognitive dysfunction along with changes in brain morphology^23,29,44,45^. However, our study utilized this model to investigate the mechanisms that link living in a low resource environment during childhood to changes in autonomic activity, BP regulation, and organ inflammation. To validate the LBN model, we recorded and analyzed the dams’ behavior during LBN exposure. We found changes in licking and grooming, self-grooming, nursing, and time off-nest, which correlated with changes that others have observed with this model^29,46^. Moreover, we calculated an entropy score, which measures maternal behavior fragmentation, and showed that our score was higher than the CONs, consistent with findings from others who use this model have found^29^ (**Table 1**).

We observed sex differences in adult BP, with LBN males showing an increase, and LBN females displaying no changes. Potential mechanisms that could facilitate these sex differences in BP are alterations in autonomic activity, inflammation, oxidative stress, vascular dysfunction, HPA axis dysfunction, endothelin-1, and decreased NO bioavailability. However, in this study, we chose to focus on autonomic activity and inflammation in the kidney and brain using the LBN model of ELS, due to their crucial role in BP regulation.

Alterations in autonomic activity, both sympathetic and parasympathetic, are known to influence BP ^47,48^. Whereas increased SNA and/or decreased PNA will elevate BP^49^. In our study, LBN males had an increase in SNA, while LBN females showed an increase in both SNA and PNA. An increase in SNA is demonstrated in both human and animal studies when the subjects are exposed to an ACE or ELS^13,43,50,51^. For example, a study investigating the association between childhood trauma and catecholamine response to psychological stressors found that 3-methoxy-4-hydroxy-phenylglycol (MHPG), a metabolite of norepinephrine, was elevated in participants who experienced ACEs^52–54^. An increase in norepinephrine concentrations in most studies suggests an increase in SNA, which is what we observed in our LBN rodents. A study by Renard *et al.* 2005 similarly found sex differences in SNA in mice exposed to maternal separation. In this study, maternally deprived females exhibited slightly higher basal plasma norepinephrine levels compared to CON females. However, plasma norepinephrine was unchanged in maternally deprived males compared to the CON^54^. Another study by González-Pardo *et al.* in 2020 further suggested sex differences in SNA, showing that norepinephrine turnover in maternally separated male mice was decreased compared to the CON, while maternally separated female mice was increased compared to CON^51^. Note that an increase in norepinephrine turnover also suggests an increase in SNA. Although these previous studies suggest some sex differences in autonomic activity with ELS exposure, our study showed that both LBN males (hypertensive) and LBN females (normotensive) had increased SNA, with females having a greater increase in SNA, which appears to be supported by clinical data. However, despite the increase in SNA in our study, LBN females also demonstrated an increase in PNA. PNA can influence BP and often works in antagonism to offset an increase in SNA^53,54^. Based on our observations, we predict that perhaps LBN male rats had increased BP due to an increase in SNA, whereas LBN female rats were protected against BP elevation due to an increase in PNA to balance out the increase SNA.

Human and animal subjects exposed to ACEs and/or ELS are associated with an increase in systemic and targeted organ inflammation throughout life^9,51,55^. An increase in inflammation is often characterized by an increase in pro-inflammatory cytokines, decrease in anti-inflammatory cytokines, and change in both types of cytokines to favor an inflammatory state^56^. The pro-inflammatory cytokines that have been augmented in human (ACE) and animal (ELS) models are TNF-α, IL-17, IL-6, and C-reactive protein^55,57^. However, these changes in pro-inflammatory cytokines are controversial and can differ depending on the species, age, and ELS events, intensity, and duration.

Two important pro-inflammatory cytokines that we investigated in the brains and kidneys are IL-17 and TNF-α, which both directly and indirectly influence BP regulation and can cause hypertension^20,58–61^. In this study, we hypothesized that both IL-17 and TNF-α would be increased in the brain and kidneys of hypertensive LBN males and not changed in normotensive LBN females. While our hypothesis was accurate with LBN females, the LBN males displayed the opposite of what we predicted. LBN males showed a reduction in pro-inflammatory cytokines (IL-17 and TNF-α) despite having elevated BP and increased SNA. Therefore, the decrease in brain and kidney pro-inflammatory cytokines may be an initial compensatory mechanism acting in response to elevated BP and/or SNA. This compensatory relationship observed in this study is not uncommon. Others have observed these same changes in pro-inflammatory markers (e.g., TNF-α and IL-1β) due to increased SNA or norepinephrine concentrations^62,63^. Thus, the changes in organ inflammation in LBN rodents may be an attempt to lower and/or maintain normal BP and to prevent tissue damage.

In summary, our data display sex differences, in which adult LBN males have elevated BP, possibly due to increased SNA. On the other hand, our data suggest that LBN females may be protected from increased BP due to a simultaneous increase in PNS and SNA. Furthermore, we detected no changes in pro-inflammatory cytokines in the brain and kidneys of LBN females but found a decrease in LBN males. The reduction in pro-inflammatory cytokines, IL-17 and TNF-α, in LBN males may serve as a compensatory mechanism to lower BP and/or prevent tissue damage. Identifying the sex differences and alterations in the pathology of adults exposed to an ELS may help to derive novel treatments for patients who have experienced ACEs.

### Limitations and Future Directions

It is important to note that this study is observational and not causal. Although the findings in this study are novel, we cannot reveal the exact timing, order of sequence of physiological events, and/or how each physiological change influences the other changes, i.e. how changes in SNA or inflammation can alter BP. To address these issues, longitudinal studies (before puberty, after puberty, and early adulthood) and gain of function and/or inhibition studies with the autonomic nervous system and/or inflammation need to be performed. Furthermore, other pro-inflammatory (IL-6, IL-1β and C-reactive protein) and anti-inflammatory (IL-4 and IL-10) cytokines can be explored in specific brain regions (such as the rostral ventrolateral medulla, paraventricular nucleus, and sections of the hippocampus) that are known to modulate BP, along with direct measurements of autonomic activity.

### Societal Implications

Understanding the mechanism of hypertension in animals that experience ELS will help scientists and physicians better understand the pathology and pathogenesis of hypertension in individuals who have experienced poverty as an ACE. Poverty is a pervasive and prevalent issue that continues to impact the health and well-being of many individuals and families worldwide. Unfortunately, with the lag in wage growth and increase in income inequality, along with a rise in inflation, food costs, high unemployment, and lack of affordable housing, it is predicted that childhood poverty rates will increase^64^. To change the negative trajectory of poor health outcomes in people that have experienced ACEs, we advocate that 1) scientists conduct more experiments to determine the impact of resource deprivation (i.e., poverty) on the development of hypertension with sex differences, and 2) the mechanisms that link ELS/ACEs to hypertension development with sex differences.

### Perspectives

The LBN model of ELS induced high BP, increased SNA, and decreased pro-inflammatory cytokines in the brain and kidneys of LBN males. However, in LBN females there was a simultaneous increase in SNA and PNA, with no changes in brain and kidney pro-inflammatory cytokines and BP. These sex difference provide knowledge and awareness into the pathology and development of hypertension caused by resource deprivation during early life. Scientists and medical providers should consider sex, the inactivation (SNA) or activation (PNA) of the autonomic nervous system, and/or organ-specific immunotherapies to assist in lowering BP, organ damage, and the risk of cardiovascular disease in adults who experienced ACEs, such as poverty.

### Novelty and Relevance

1. **What Is New?**

- LBN exposure increases BP in males, but not females, possibly due to alterations in autonomic nervous system activity.
- The reduction in localized pro-inflammatory cytokines may be a compensatory mechanism to lower BP and/or protect from tissue damage in LBN males.
2. **What Is Relevant?**

- Childhood poverty is a major public health concern that increases the risk of hypertension later in life, which presents differently in males and females.
- Using the LBN model helps to elucidate the sex differences and mechanisms connecting childhood poverty to hypertension.
3. **Clinical/Pathophysiological Implications?**

- This study helps uncover potential therapeutic targets and bring awareness to the sex differences in hypertension in individuals who experienced childhood poverty.

## Abbreviations

(ACE): Adverse childhood experience
(A.U.): Arbitrary Units
(BP): Blood pressure
(BP): Centers for Disease Control
(BP): Early life stress
(ELISAs): Enzyme-linked immunosorbent assays
(FFT): Fast Fourier transform
(GD): Gestational Day
(HR): Heart Rate
(HFHRV): High frequency heart rate variability
(HPA): Hypothalamic-pituitary-adrenal
(IACUC): Institutional Animal Care and Use Committee
(IL-17): Interleukin-17
(LSD): Least significant difference
(LFBPV): Low frequency blood pressure variability
(NIH): National Institutes of Health
(PND): Postnatal day
(RPM): Revolutions per minute
(SEM): Standard error of the mean
(TNF-α): Tumor necrosis factor
(UNTHFW): University of North Texas Health Fort Worth

## Acknowledgements

The authors acknowledge the support of the Department of Physiology and Anatomy at UNTHFW.

## Availability of Data

The data that support the findings of this study are available from the corresponding author, [Mark Cunningham], upon reasonable request.

## Author Contributions

**Jonna Smith** (***first author of the manuscript***): formal analysis, experimental investigation, methodology, project administration, data curation, visualization, writing-original draft, writing-edited and revised manuscript. **Savanna Smith, Kylie Jones, Angie Castillo, Faith Femi-Ogunyemi, and Allison Burkes:** experimental investigation, data curation, methodology, resources, writing-edited and revised manuscript. **Jessica Bolton, Ahfiya Howard, and Luis Colon-Perez:** writing-edited and revised manuscript. **Mark Cunningham** (***PI***): conceptualization, formal analysis, funding acquisition, investigation, methodology, project administration, resources, supervision, validation, writing-review and editing, helped to perform experiments, and approved final version of manuscript.

---

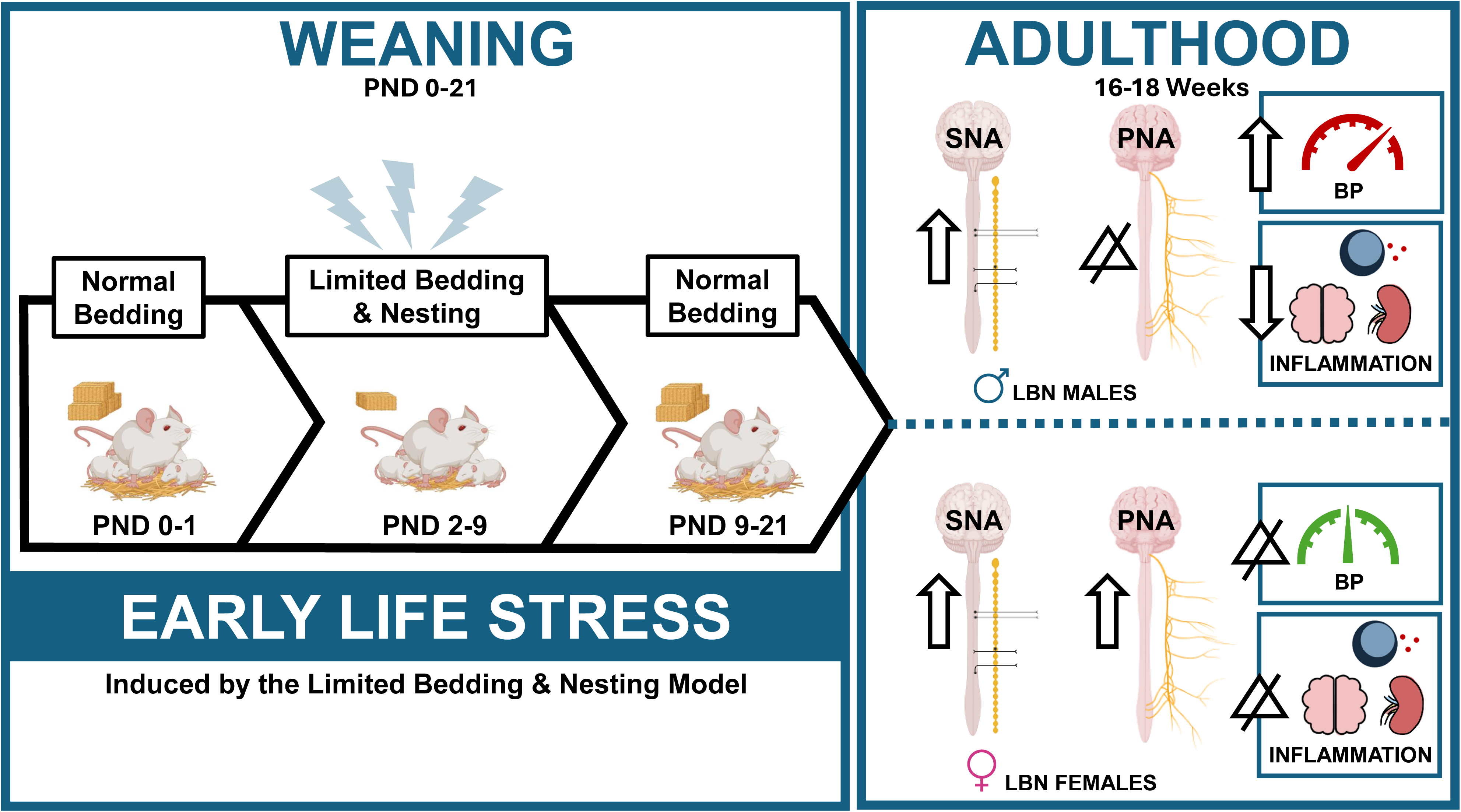

